# Reversibly-bonded Microfluidic Devices for Stable Cell Culture and Rapid, Gentle Cell Extraction

**DOI:** 10.1101/2023.12.06.570496

**Authors:** Xiaohan Feng, Lily Kwan Wai Cheng, Xuyan Lin, Angela Ruohao Wu

## Abstract

Microfluidics chips have emerged as significant tools in cell culture due to their capacity for supporting cells to adopt more physiologically relevant morphology in 3D compared with traditional cell culture in 2D. Currently, irreversible bonding methods commonly used in chip fabrication mean that chips cannot be detached from their substrate without destroying the chip structure, which makes it challenging to do further analysis on cells that have been cultured on-chip. Some reversible bonding techniques exist but are restricted to certain materials, or require complex processing procedures. Here, we demonstrate a simple and reversible polydimethylsiloxane (PDMS)-polystyrene (PS) bonding technique that allows devices to withstand extended operation while pressurized, and supports long-term stable cell cultures. Importantly, it allows rapid and gentle live cell extraction for further downstream manipulation and characterization after long-term on-chip culturing, or even further subculturing. Our new approach could greatly facilitate microfluidic chip-based tissue and cell cultures, overcoming current analytical limitations and opening up new avenues for downstream uses of on-chip cultures, including 3D-engineered tissue structures for biomedical applications.

## Introduction

Microfluidic devices have seen extensive utilization in the domain of cellular biology.[1, 2] For instance, 3D tumor models, cell-interaction models, and organ on-chips have been successfully constructed using microfluidics for elucidating biological mechanisms and for drug discovery.[3-3] The conventional polydimethylsiloxane (PDMS) chip is usually irreversibly bonded to the substrate and therefore cannot be “opened” unless the device is destroyed. [6] This makes such devices suitable for long-term cell culture, but efficiently extracting a substantial number of viable cells from the device for downstream manipulation is challenging, even with the aid of Trypsin. Due to this challenge, analysis methods on cells grown in microfluidic devices are also limited, most of the time restricted to imaging, bulk molecular analysis, or simple on-chip staining assays.

Achieving reversible bonding between PDMS and substrate is a possible solution to this limitation, and efforts have been made in this direction. The tight irreversible bonding of devices can be easily achieved by plasma treatment and chemical modification such as 1% APTES treatment[7, 8], but long-term stable methods for reversible bonding are not so easy to achieve. Existing methods for reversible bonding work on a limited range of device-substrate material pairs, or require complex processing procedures. For instance, increasing the thickness of PDMS and bonding to glass without plasma treatment is the simplest way to create reversible bonds,[9] but this method of bonding is less robust and suffers from low-pressure tolerance, with a high risk of leakage under pressurized or prolonged usage, which makes it non-ideal for long-term cell culture or scenarios that require pressurized pumping and control. More recently, a glass-based device manufacturing approach was proposed for achieving reversible bonding and gentle cell extraction.[10] The reversible bonds in this approach are created by water dehydration between two glass slips with high cleanliness, therefore requiring neutral detergent and continued exertion of external force for forming bonds. Also, this method is not applicable to PDMS-based devices, which are widely used for cell-based studies. We are still in need of a versatile and simple method of reversible bonding that is applicable to commonly used device-substrate materials.

Another common approach to create reversible bonds is providing a sacrifice layer within devices, which can be designed with multiple materials, such as polymethylmethacrylate (PMMA), polycarbonate (PC), polystyrene (PS), and polyethylene terephthalate (PET/PETG).[11-13] For example, Thompson et al. used adhesive tape for creating reversible bonds within complex devices,[12] but the low manufacturing throughput and as yet undetermined biocompatibility limits the applicability of this method. Similarly, a silicone-based soft skin adhesive was mixed with PDMS to create a sacrifice layer between the PDMS device and their substrate for long-term cell culture,[13] but the low-adhesion property of this adhesive may influence cell attachment on devices, thereby affecting cellular biomechanics or function on-chip. Moreover, there are other strategies for creating reversible bonds such as wax-bonding,[14] reducing curing agents,[15] clamping strategies,[16, 17] changing substrate,[18] and termed sandwich bonding.[19] However, these methods often involve intricate manufacturing processes that may limit scaled production. Furthermore, using the aforementioned methods to create complex designs for multilayer or multichannel devices becomes more difficult. Most importantly, efficient live cell extraction has not yet been reported for any of these previous approaches.

In this study, we present a new approach for fabricating reversibly-bonded microfluidic devices (RBM devices, and here irreversibly-bonded microfluidics (IRBM) is corresponding to the microfluidic devices whose substrates cannot be manually separated with the cover slabs), which is achieved by applying a 0.1% (3-Aminopropyl) triethoxysilane (APTES) solution to a PS substrate. The low concentration of 0.1% APTES solution introduces a small proportion of amine groups for covalent bonding, and it roughens the PS surface thereby enhancing van der Waals forces, overall resulting in a stable and reversible bonding between PS and PDMS. The procedure requires no other equipment beyond a benchtop plasma cleaner, and can be done outside the cleanroom, making it accessible to most laboratories that make PDMS devices and is compatible with most PDMS-based device fabrication workflows. We demonstrate the biocompatibility of this type of reversibly bonded device by performing long-term and stable cell culture on-chip and achieving high cell viability, as well as show tumor spheroid formation on-chip as an additional application.

We additionally illustrate the biocompatibility of this surface treatment using vascular cell culture and vascular network formation experiments on-chip. Next, we recover cells from the chip by rapid hand-peeling of the PDMS slab and verify the high efficiency of live cell extraction by live cell counting, cell recovery experiments, and flow cytometry. Finally, we use a device design with multiple compartments to co-culture different cell types, and show that the reversibly bonded device can be used for compartmentalized cell separation with minimal cross-contamination, which further extends the versatility of this bonding method. Overall, this reversible bonding method offers a robust and scalable chip fabrication process, has high biocompatibility, and allows for gentle cell extraction from microfluidic devices. The viable cells extracted from reversible devices can contribute to further understanding of cellular behaviors and the mechanisms behind them within the engineered microenvironment.

## Experimental

### Microfluidic device design and fabrication

The microfluidic devices used in this study were made of PDMS and modified from a previous study.[20] Soft lithography and replica molding were used to fabricate the device. Briefly, the lithography mask was designed using AutoCAD. A silicon wafer was then patterned by SU8-2050 photoresists (Microchem Corp., Westborough, USA) using the standard photolithography techniques (ABM/8/500/NUV/DCCD/SA, ABM, USA). PDMS (Sylgard 184 Silicone Elastomer, Dow Corning, USA) was mixed at a 10:1 ratio of elastomer to curing agent and then poured over the silicon mold. After degassing, the PDMS was cured on the mold for at least 2 hours at 80 °C, until fully cured, before peeling off. The inlets and outlets of the PDMS chambers were punched using a PDMS puncher prior to substrate bonding. A petri dish made of PS (#150318, ThermoFisher, USA) was used as the substrate for bonding, and it was treated with oxygen plasma discharge in a plasma cleaner (PDC-002, Harrick Plasma, New York, USA) for 3 minutes, followed by coating with 0.1% v/v APTES ethanol dilution for 10 minutes at ambient temperature. Afterward, the dish was washed with Dulbecco’s phosphate-buffered saline (DPBS, 14190235, ThermoFisher) until the substrate surface became hydrophilic. To completely remove the residual liquid, the cleaned PS dishes were placed on a hotplate at 75°C for primary drying for 1 to 2 hours before fully drying out using a nitrogen gas spray gun. The PDMS slab to be bonded was cleaned with IPA, distilled water, and scotch tape, and dried with an air spray gun. To form the reversible bonding between the PDMS slab and PS substrate, both the PDMS and the PS petri dish were treated with oxygen plasma for 3 minutes, followed by immediately placing the PDMS slab surface in contact with the PS dish surface, and making sure a good seal has formed between the two surfaces by pressing. Finally, the bonded device was incubated overnight on a hotplate at 75°C and sterilized by UV irradiation for at least 30 minutes before use.

### Surface characterization

The PS substrate was cut into small pieces, sputter coated, and mounted, and then subject to materials characterization of its functionalized surface. We used surface electron microscope (SEM) (JSM-6320F, JEOL, JP), X-ray photoelectron spectroscopy (XPS) (PHI 5000 Versaprobe III, ULVAC-PHI, Japan), and atomic force microscopy (AFM) (Dimension ICON, Bruker, MA, USA) to characterize the surface properties. The XPS analyses were conducted using a machine equipped with an aluminum X-ray source (mono-gun, 1486.6 eV) with a pass energy of 40 eV. The binding energy of C 1s (284.5 eV) was used as the reference. The resolution for the measurement of the binding energy was approximately 0.1 eV.

### Cell culture

A human glioblastoma cell line (U87) was used in cell viability and tumor spheroid formation experiments. A human hepatocellular cell line (HepG2) was used in cell extraction and separation experiments. Both were cultured in Dulbecco’s Modified Eagle Medium (DMEM, 11965-092, ThermoFisher) with 10% fetal bovine serum (FBS, A4766801, ThermoFisher). Human umbilical vein endothelial cells (ECs, CC-2519, Lonza) were used for the vascular network formation experiments, and were cultured in endothelial microvasculature growth medium (EGM-2MV BulletKit™, CC-3202, Lonza); normal human lung fibroblasts (FBs) (NHLFs, CC-2512, Lonza) were cultured in fibroblast growth medium (FGM-2 BulletKit™, CC-3132, Lonza). To support vasculature forming in the device, FBs were transferred to the EGM-2 MV medium and cultured in it for at least 2 passages before proceeding on-chip co-culture with HUVECs. Monocyte cell line THP-1 was used in cell separation experiments and cultured in RMPI-1640 (11875119, ThermoFisher) with 10% FBS. Cells were cultured on Petri dishes under a humidified incubator at 37 °C and 5% CO_2_ and grown up to 80% confluency for cell seeding experiments.

### Gel preparation and cell seeding

Cell seeding gel preparation was performed as previously described.[20] To generate the fibrin gel, the Fibrinogen solution and thrombin solution were prepared separately. Briefly, the fibrinogen solution was prepared by dissolving 15 mg of bovine fibrinogen (F8630-1G, Sigama-Aldrich) in 2.5 ml of DPBS in a 37 °C water bath for 1 hour, then further sterilized by filtered using a 0.22μm filter. The thrombin stock solution was prepared by dissolving thrombin (T9549, Sigma-Aldrich) with 0.1% w/v BSA solution (B14, ThermoFIsher) into the concentration of 100 U/ml, which was then stored in -20 °C. Before use, the thrombin solution was diluted into 4 U/ml by adding the culture medium. For cell seeding, U87s, ECs, FBs, and HepG2s were trypsinized using TrypLE (#12605028, ThermoFisher) for 5 minutes, neutralized with culture medium, and then centrifuged for 3 minutes at 300g. After aspirating the medium off, the cell pellets were resuspended with diluted thrombin solution, and the cell solution were then quickly mixed with filtered fibrinogen solution. The cell-laden gel mixtures (3 or 6 million cells per ml of U87s for cell viability experiments; 3 million cells per ml of Ecs and 1.5 million cells per ml of FBs for forming vascular networks on-chip; 3 million cells per ml of HepG2s for cell extraction and separation experiments) were then gently introduced into individual channels and allowed to polymerize for 15 minutes at room temperature by putting them into a humidified chamber. The corresponding channels were filled with cell culture medium when those gel mixtures were cross-linked After that, the channels for the culture medium were loaded by pipetting. All the devices were incubated under a humidified incubator at 37 °C and 5% CO2, and the cell culture medium was removed and refilled with fresh culture medium every 24 hours.

### Cell viability calculation

The cell viability was calculated by the formula below with separately counted live and dead cells through ImageJ. The LIVE/DEAD Cell Imaging Kit (488/570) (#R37601, Thermofisher, USA) was applied to stain the live and dead cells separately for confocal imaging.

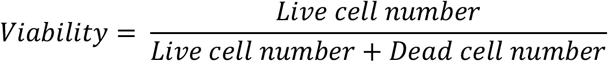

### Cell extraction and separation

The microfluidic chip containing cells is carefully removed from the incubator before adding the TrypLE to the chip. For the IRBM chip, approximately 400 μL TrypLE is added to the chip surface and the channel near to cells (Fig. 1A). For the RBM chip, approximately 200 μL of TrypLE is added to both the substrate and the PDMS chip after the PDMS slab is manually peeled off from the underlying substrate (Fig. 1B). Then, an additional 200 μL of culture medium is added to the chip after incubating at 37°C, and the neutralized cell suspension is subsequently collected (Fig. 1A-B). The live cell counting was achieved by Countess II (#AMQAX1000, Thermofisher, USA) after adding Trypan Blue (#15250061, ThermoFisher, USA) to the cell suspension. The fibrin gel is utilized for location-specific cell separation. During the cell separation process, it is imperative to treat the substrate and PDMS chip with individual processing steps. The separation ratio (SR) of a specific region was calculated by the formula below:

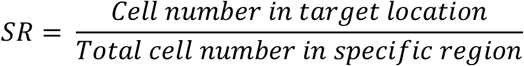

**Fig. 1.**
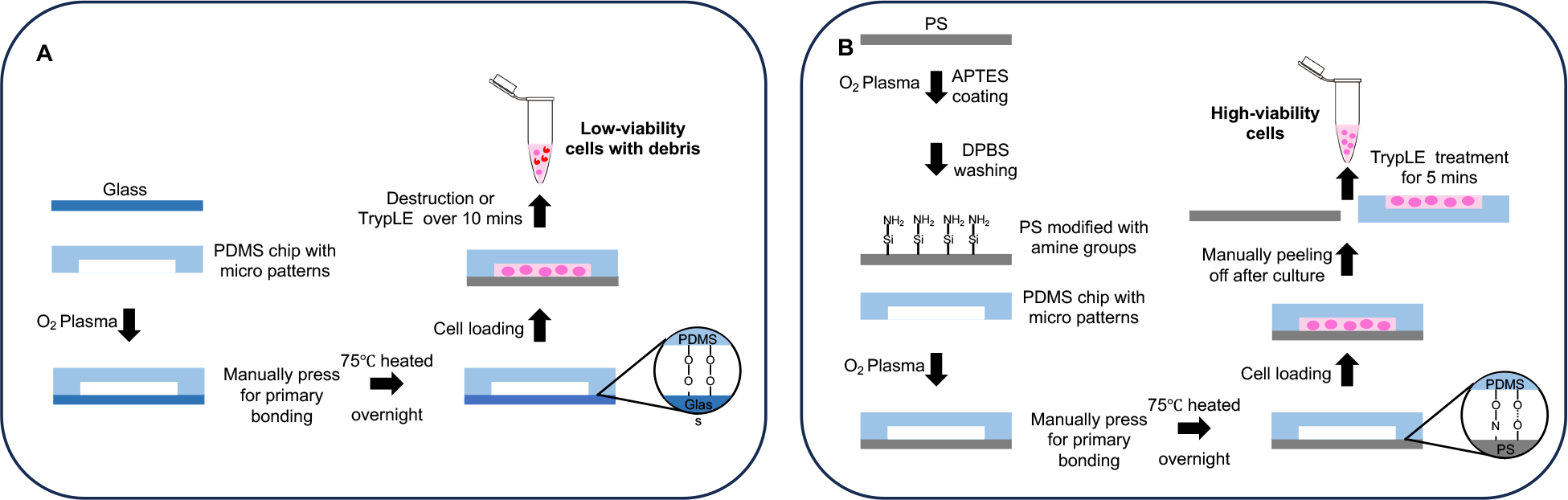
A: Cell extraction workflow of conventional irreversible bonding devices; B: Process flow of reversible bonding devices and rapid cell extraction

For instance, in calculating the separation ratio of the region with fibrin gel, the “cell number in target location” refers to cells on the PDMS chip, while the “total cell number in the specific region” encompasses cells on both the PDMS chip and the substrate.

### Burst pressure test

The bond strength of the interface was measured *via* a burst pressure test for PDMS to glass substrate after plasma treatment (IRBM control) and PDMS to APTES-treated PS dishes (RBM devices). For each test, the PDMS slab that had a 2 mm length, 1 mm width, and 50 μm height chamber was pressurized incrementally until bursting between PDMS and substrate occurred. The burst pressure test setup consisted of a closed system in which the tubing was connected to an air pump with a digital manometer to the microfluidic chamber. Measurements were conducted after the pressure reached equilibrium, and the final stable pressure, just prior to any PDMS-substrate burst, was documented for each device.

## Results and discussion

### Reduced APTES concentration facilitates reversible bonding between PS and PDMS

Oxygen plasma treatment is a typical and conventional bonding method for PDMS-based devices, enabling the irreversible bonding of PDMS-glass chips by activating the surface siloxane bonds of PDMS. Additionally, oxygen plasma followed by chemical treatment was also applied to other materials to enhance the bonding strength of PDMS-based devices.^[21]^ For instance, plasma-activated PDMS can bond to a PS substrate with a weak bonding strength of around 1.72 psi. However, irreversible bonding can be achieved by treating the PS substrate with 1% APTES solution after oxygen plasma treatment.^[22, 23]^

To create a reversibly bonded device suitable for long-term cell culture and cell extraction, we used 0.1% APTES ethanol solution to treat both PS and PDMS. This treatment resulted in reversible bonding between the two surfaces, enabling long-term cell culture in the PDMS device, with the added feature that the bonded surfaces can be easily separated by hand-peeling.

SEM analysis was utilized to explore the bonding mechanism between PDMS and 0.1% v/v APTES treated-PS. The analysis confirmed the anticipated adhesion, which is shown in Fig. 2A-C. Fig. 2A showed the untreated PS surface before the APTES coating, and Fig. 2B displayed the PS surface after being treated with APTES. According to SEM images, the untreated or treated PS surface maintained a smooth surface regardless of the APTES treatment, suggesting there was no significant morphology change on the PS surface after oxygen plasma and low-concentration APTES treatment (Fig. 2A-B). After peeling the PDMS slab off from the PS substrate, a few silicon residues along with partial deformation of the substrate were observed (Fig. 2C). These observations suggest the existence of covalent bonds between the PDMS and the treated PS, which have been induced by oxygen plasma treatment and APTES treatment. The substrate deformation can be attributed to the delamination process, suggesting that chemical bonding occurs between PDMS and treated PS, rather than a simple stacking of the two materials. This bonding strategy evidently differs from the conventional PDMS-glass or APTES-treated irreversible chips, as there are only small silicone fragments remaining on the substrate after peeling, indicating the limited amount of chemical bonding between PDMS and treated PS results in the desired stable and reversible bonding for this device.

**Fig. 2.**
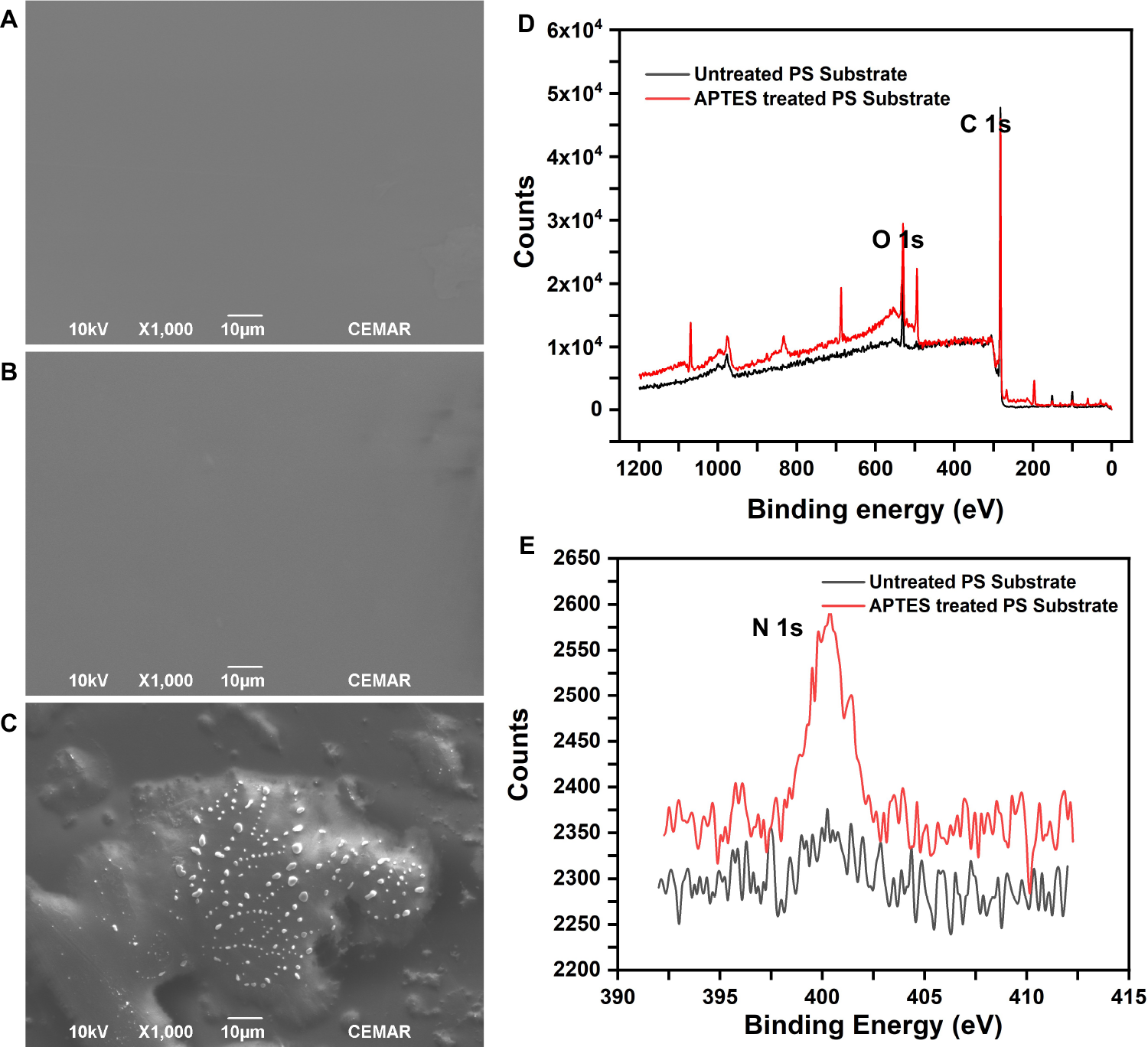
A: The SEM image of untreated PS surface; B: PS surface after treatment with a 0.1% APTES solution; C: PS surface after peeling off the PDMS; D: XPS survey spectrum for untreated PS and APTES-treated PS; E: N 1s fine spectrum, which indicates amine groups on treated PS surface.

The effects of the 0.1% v/v APTES solution treatment are elucidated through the XPS full spectrum and Nitrogen (N 1s) fine spectrum, as depicted in Figures 2D and 2E. By comparing the full spectrum of untreated PS with APTES-treated PS, additional peaks were found in the spectrum of APTES-treated PS, (Fig. 2D), indicating the presence of additional chemical groups at the PS surface after treating with 0.1% v/v APTES solution (Fig. 2D). The appearance of new peaks at 400 eV corresponds to the N 1s spectra (Fig. 2D-E), indicating the successful incorporation of amino silane onto the PS surface. The existence of amino functional groups may form N-O bonds with the PDMS slab after plasma treatment, which contributes to the bonding strength. However, the counts of N 1s peak on the PS surface after 0.1% APTES solution treatment was significantly lower than that of PS treated with 1% APTES solution treatment,[7] which demonstrates that a limited amount of covalent bonding resulted from low-concentration APTES treatment, and that this lowered capability for covalent bonds formation is crucial for reversible bonding.

### Surface morphology reveals the key of reversible bonding is water-weakened bonds

Having illustrated the effect of low-concentration APTES treatment on the PS surface, we sought to quantify the bonding strength and further explore the underlying mechanism responsible for the bonding strength variation when a lower concentration of APTES is used.

We conducted two different pressure tests between conventional IRBM chips and RBM chips to demonstrate the bonding strength of the latter. In gas pressure tests, both types of chips showed comparable performance: both can withstand a gas pressure of up to 14.5 psi without any bursting (Fig. 3A). In fluid pressure tests, the pass rate for the RBM chips was 66.7% at 12 psi, while the pass rate for the corresponding IRBM chips remained at 100%. Here the pass rate is defined as the proportion of chips that operate under this pressure without failure for at least half an hour. Upon increasing liquid pressure to 14.5 psi, the pass rate for reversible bonding chips dropped to 55.6%. The reduction in pass rates under liquid pressure is likely due to the infiltration of water molecules to the PDMS-PS interface, which weakens the bonding strength. To understand the mechanism for this change in the bonding strength after the device has been used with aqueous reagents, we used AFM analysis to check for morphology transformations on surfaces under different conditions (Fig. 3B-E). A relatively flat surface profile was exhibited by the untreated PS surface (Fig. 3B). After treating the PS surface with APTES and prior to bonding, we saw a significant increase in PS surface roughness. The increase in surface roughness elevates the substrate’s surface free energy, thereby enhancing its bonding characteristics when dry. We also noted that manual delamination of PDMS without prior water treatment led to substantial silicon residue on the PS surface (Fig. 3D). We performed the measurement again with APTES-treated PS that was bonded, treated with water to mimic the device contact with aqueous reagents such as buffer and media, and subsequently manually delaminated. This time the PS surface had less remaining silicon residue after delamination compared with the sample that did not undergo water treatment, and also had a reduced surface roughness (Fig. 3E). These results indicate that contact with water disrupts the bonding between the PDMS slab and PS substrate, thus easing the PDMS peeling process after cell culture. Overall, this water-weakened bonding strength is the basis of the reversible technique and makes hand peeling feasible.

**Fig. 3.**
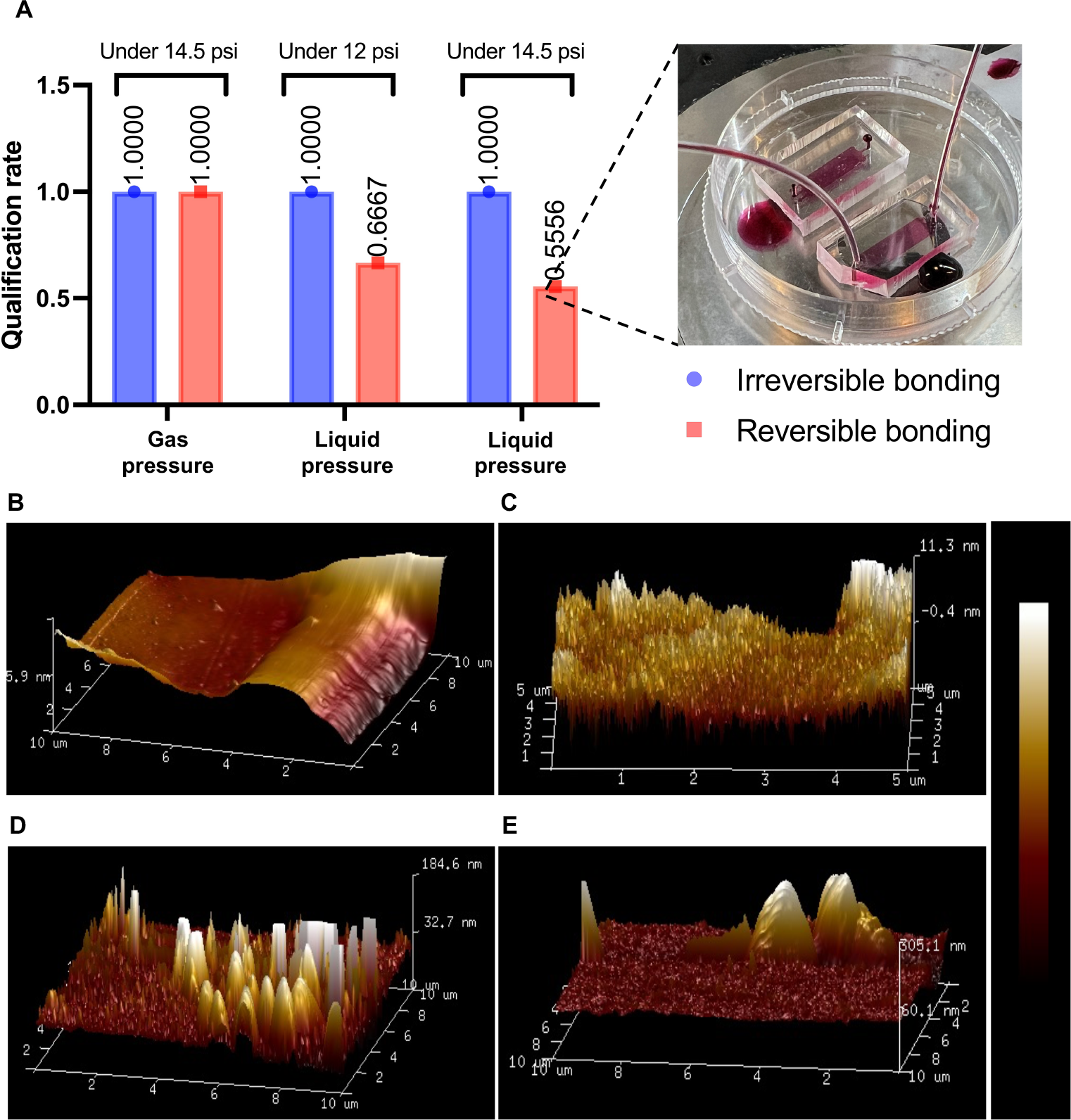
A: Pressure test results for conventional IRBM chips and RBM chips (n=9), both reversible and irreversible devices have 100% pass rates under 14.5 psi gas pressure (maximum pressure of testing machine), and pass rates of reversible devices under 12 psi and 14.5 psi liquid pressure are 66.7% and 55.6% respectively; B: AFM images of untreated PS; C: APTES-treated PS; D: PS by manual delamination without water treatment; E: PS by manual delamination treating with water.

### Reversible devices demonstrate high biocompatibility by culturing multi-types of cells

Reversible bonding is essential for recovering cells from microfluidic devices for further analysis.[23] Another basic requirement of RBM devices is biocompatibility, so that cells can live, propagate, and maintain their normal functions.

After testing the physical performance on the reversible device, we next characterized the feasibility of this RBM device for cell culture. The U87 glioblastoma cell line was used to determine the biocompatibility of reversible chips and calculate cell viability. After cell seeding on-chip, the U87 cells exhibited appropriate epithelial morphology after culturing on-chip for a few days, and even formed spheroids when culture period was extended, which indicates their robust growth on the chip (Fig. 4A-C). On D1, D4, and D7, Live/Dead cell kits were applied to observe the growth status of U87 on the chip (Fig. 4A). At those selected time points, using confocal microscopy, we observed a large amount of green fluorescence indicating live cells, as well as a small amount of red fluorescence indicating dead cells. The calculated cell viabilities at selected time points are shown in Fig. 4B. The cell viability at D4 and D7 was nearly 100%, which indicates that the glioblastoma cells grew well on the RBM device. On D1, there was ∼80% on-chip cell viability due to cell loss during passaging.

**Fig. 4.**
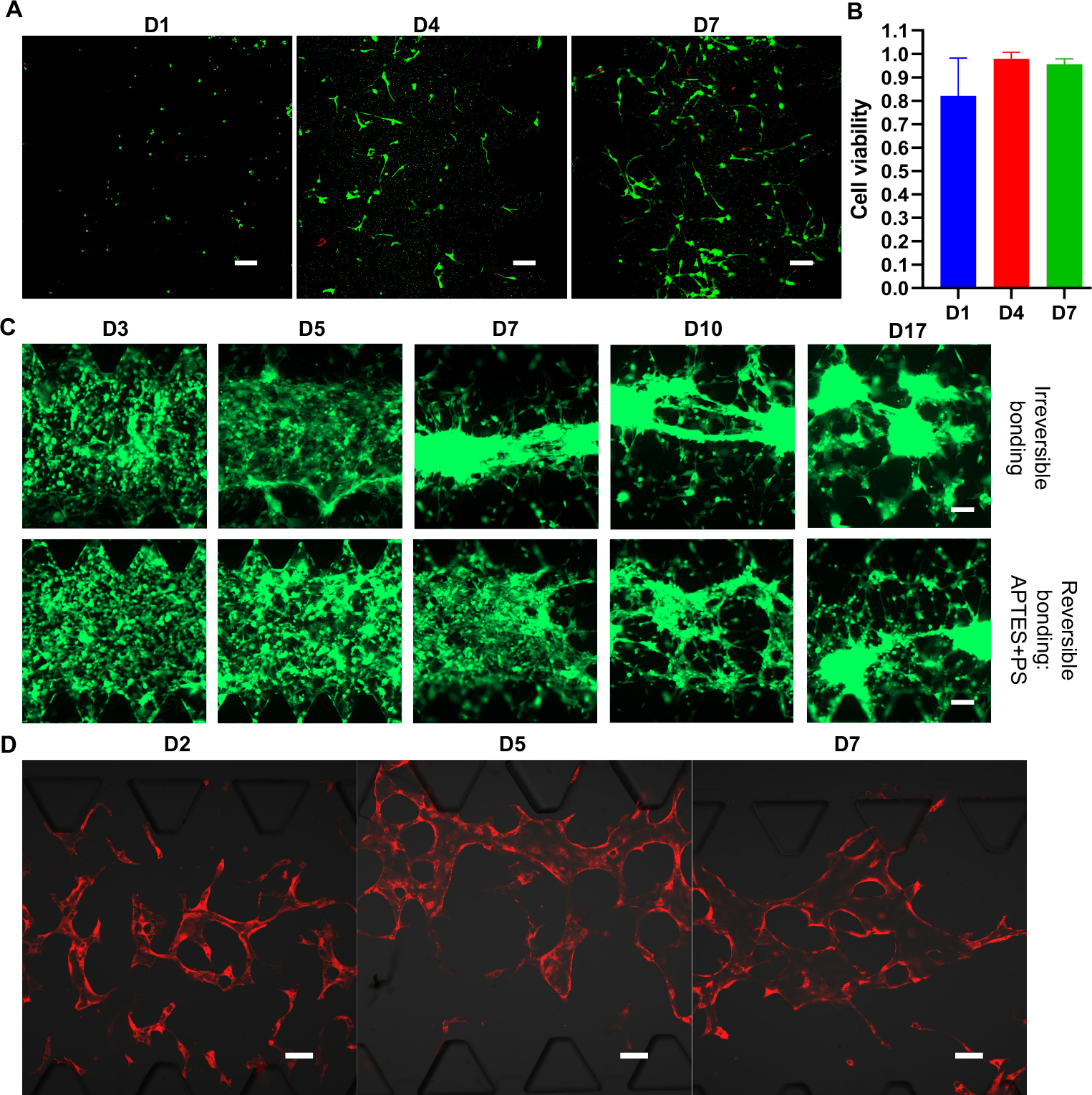
A: Confocal images after treating U87 on-chip with Live/Dead cell kit, the U87 cells begin to show epithelial morphology at Day 4; B: Cell viability calculated by ImageJ processing on confocal images at different time points; C: U87 spheroid formed on RBM chip at the later stage of culture; D: Confocal images of vasculature on RBM device at different timepoints (bar=100μm).

Next, we tried to increase the seeding concentration of U87 cells and extend the on-chip culture period to more than 15 days. The U87 cells could form tumor spheroids and extend to form cell-cell connections between spheroids regardless of the bonding approach (Fig. 4C). The formation of spheroids on RBM chips lags slightly behind that of IRBM chips, which could be due to the lowered cell migration associated with different surface roughness of PS substrate with and without APTES treatment.

Tumor cells are generally considered to be more robust in culture than other types of differentiated cells or primary cells. Hence, primary HUVECs and FBs were cultured in RBM devices to further assess the biocompatibility of this device with more delicate cell types. We seeded the HUVECs and FBs in different channels at their designated concentration, as previously described.[20] Notably, the HUVECs could form lumen-like structures after 5 days of on-chip culture (Fig. 4D), showing that reversible devices not only do not negatively affect the growth of HUVECs but also allow them to form functional lumen-like structures. In general, all seeded cells displayed expected growth rates on the reversible devices regardless of cell type, which validates the high biocompatibility of this RBM device.

### RBM devices enable rapid and gentle cell extraction and separation

In addition to biocompatibility, the efficient and gentle extraction of cells from the device is also crucial for downstream cell profiling, such as FACS analysis, sorting and subculturing, and genomics sequencing. A gentle cell retrieval process is necessary to minimize cell loss and avoid dramatically altering the cell state. We next demonstrated a simple method of retrieval of cells cultured in this RBM device, and assessed the damage to the cells during extraction.

The HepG2-mCherry liver tumor cells were seeded and cultured for 5 days on-chip. Before cell retrieval, confocal images were taken of cells seeded and cultured on-chip, shown in Fig. 5A. After cell retrieval, the collected cell suspensions were re-seeded onto a conventional culture dish for cell recovery, and confocal images were taken after six days of conventional culturing on-dish. For IRBM chips, trypsin was pushed into the device to detach cells from the fibrin gel and flushed out for collection, whereas for RBM chips, the PDMS was peeled off by hand, and then the cell-embedded side was subject to trypsinization. Peeling off allows trypsin to efficiently reach and treat the gel-embedded cells, and allows thorough washing to collect the detached cells as much as possible. After retrieval, only the cells recovered from RBM devices showed fluorescent signals, indicating successful cell subculturing (Fig. 5B). Additionally, partial cell suspensions were subject to FACS to calculate live cell numbers: 1448 ± 1698 and 5289 ± 1596 for six IRBM and RBM devices respectively (Fig. 5C). These results suggest that only very few cells could be successfully extracted from the IRBM chip by flushing, while our RBM chip allowed extraction of many more cells, with an estimated extraction rate of 19.2% and 55.6%, respectively. The difference in cell recovery results and extraction efficiencies between IRBM and RBM devices indicates the excellent cell extraction performance of RBM devices.

**Fig. 5.**
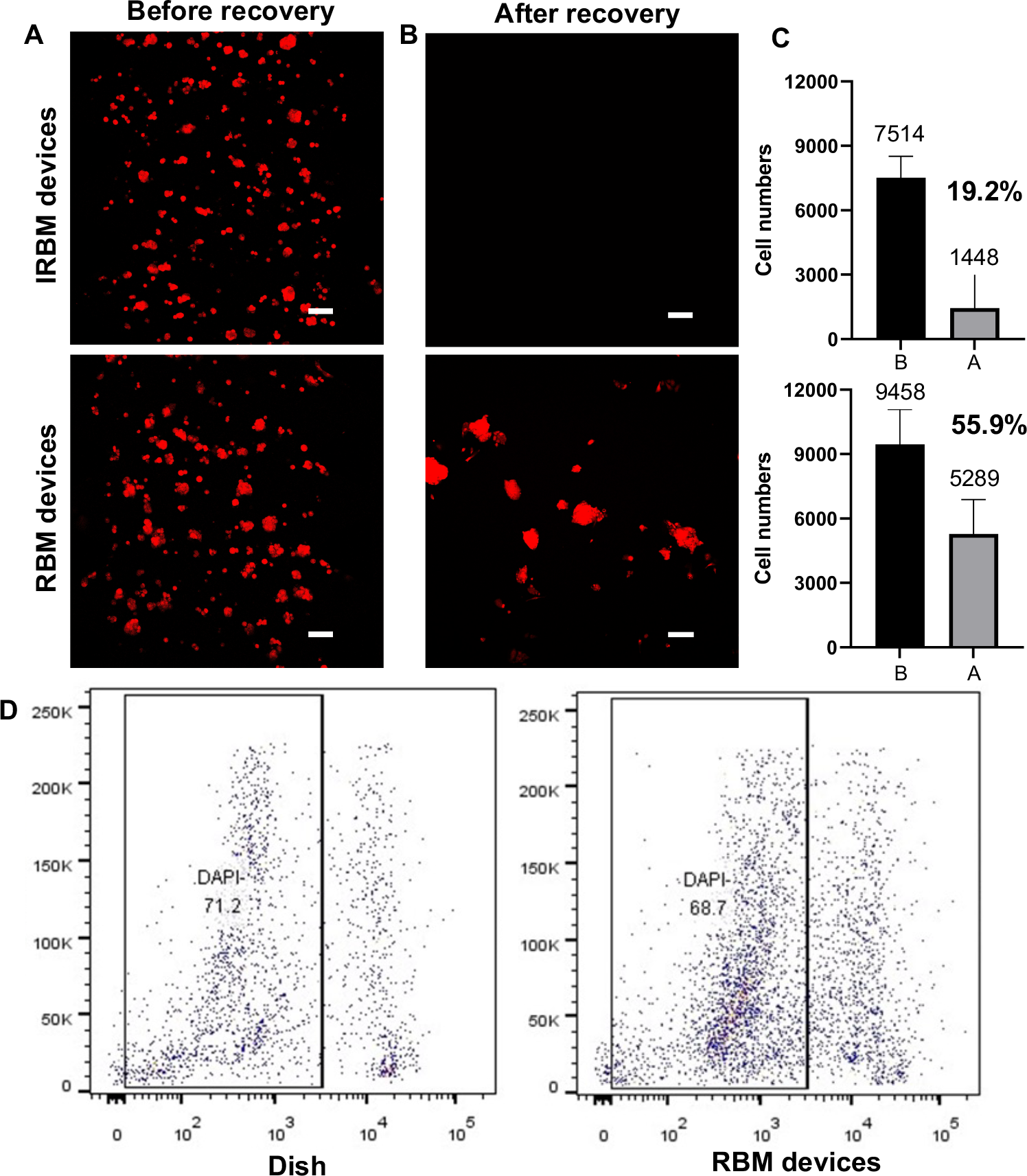
A: Confocal image of HepG2 before cell extraction on IRBM and RBM chip (bar=100μm); B: The confocal image of HepG2 cell recovery culture for 6 days on IRBM and RBM chip (bar=50μm); C: Extraction efficiency of live cells from IRBM and RBM chip (n=6, black and grey bars represent cell numbers of before extraction and after extraction respectively); D: Flow cytometry results indicate that extracted cells’ viability from culture dish and RBM devices.

Furthermore, we compared the damage caused by cell retrieval from RBM devices and normal culture dishes by DAPI staining flow cytometry. The live cell percentages are similar between dish culture and RBM devices, at 71.2% and 68.7% respectively (Fig. 5D). Although the live cell percentage from the RBM chip was slightly lower than that of the traditional cell extraction method from dishes, our method can extract sufficient cells from PDMS microfluidic chips for subsequent manipulation, such as single-cell sequencing.

Apart from extracting cells, collecting cells with distinct behaviours from microfluidic chips also presents challenges to researchers, particularly those employing devices to mimic cell behaviors and tissue niches, such as T cell migration within the tumor microenvironment (TME) on-chip.[5, 24] However, the traditional extraction methods cannot achieve separate retrieval of migrated or unmigrated cells due to device destruction during extraction. Capable of both robust cell culture and efficient cell extraction, our RBM device can also be used to separately retrieve cells from different regions or chambers on the chip. The cell extraction platform was enhanced by integrating fibrin gel, leading to the development of a region-based cell separation device (Fig. 6A). Within this device, the attaching properties of compartmentalized cells are distinguished apart by the assistance of fibrin gel. Cells without fibrin gel attach to the PS substrate, and other cells are aided by the fibrin gel in attaching to the PDMS chip after manually peeling. This difference in attaching properties makes cell separation possible on this device.

**Fig. 6.**
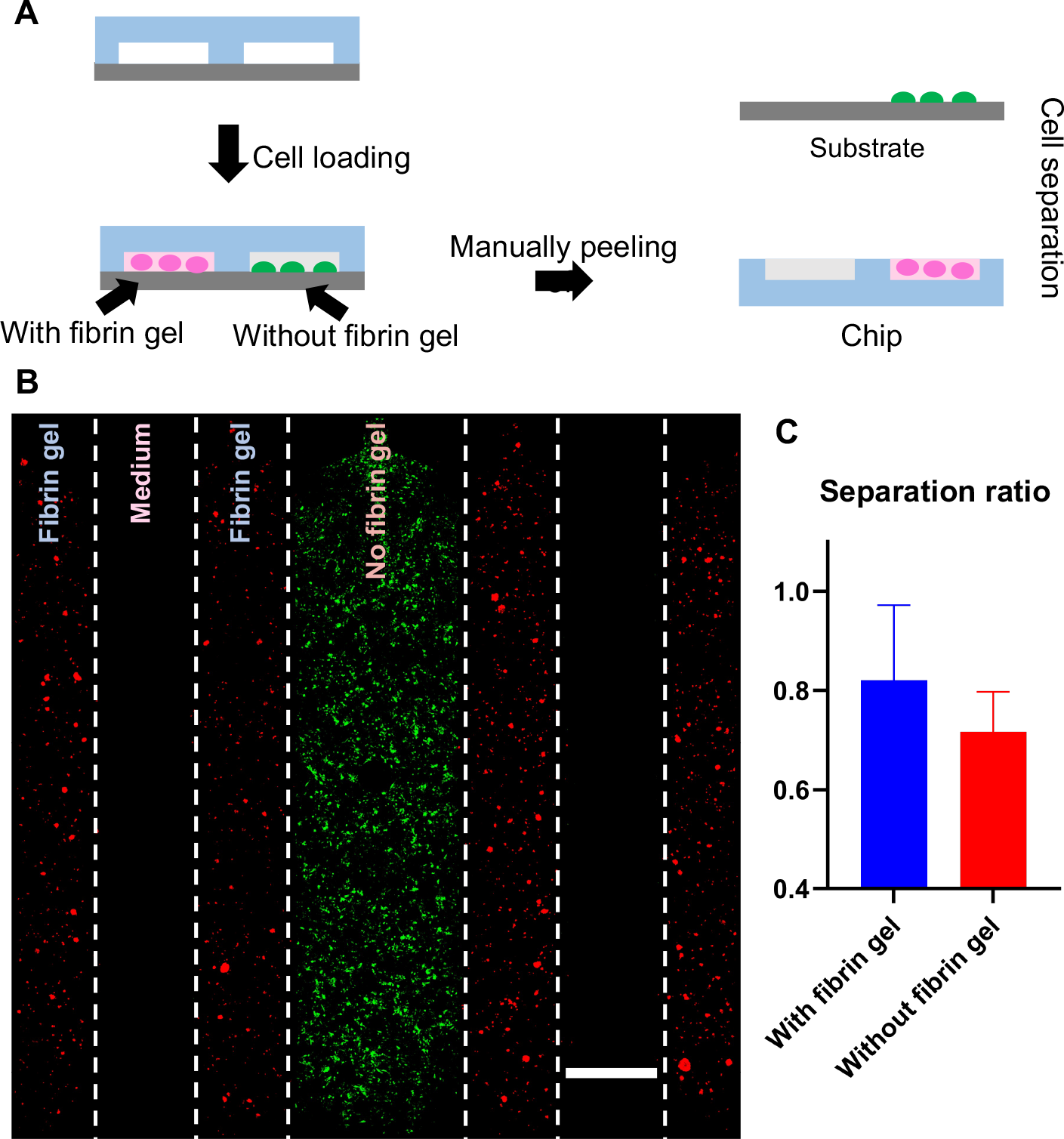
A: Workflow of cell separation from the device; B: Different cells seeded in the different regions of the multi-channel chip for mimicking cell interactions in TME; C: Above 70% separation ratio of with or without fibrin gel (n=3).

We conducted preliminary testing on a multi-channel PDMS chip, introducing THP-1 cells with green-fluorescent in the middle channel and seeding HepG2-mCherry cells on both sides to mimic a hepatic TME (Fig. 6B). The compartmentalized cells could be collected separately after peeling off the chip from the substrate. The SRs with and without fibrin gel are 82% and 72% respectively, which display acceptable ratios for separating cells (Fig. 6C). With the assistance of fibrin gel, the motility of cells will be limited and slightly increases the SR. However, these separation rates could be further improved by reducing the number of seeded cells or shortening the culture time.

Overall, the application of this reversible bonding approach enabled the microfluidic device to achieve both streamlined on-chip cell extraction and rapid compartmentalized cell separation.

## Conclusions

In this paper, we developed an approach to achieve reversible PDMS-PS bonding by treating low-concentration APTES on PS, thus overcoming many limitations of live cell retrieval from PDMS-based microfluidic cell culture devices. This RBM device can withstand pressures exceeding 12 psi in most cases, providing sufficient pressure resistance for the practical use of many applications, and enabling long-term stable cell culture. We observed high cell viability for multiple cell types, including glioblastoma cell line, hepatic cancer cell line, and endothelial primary cell line, which highlights the biocompatibility of these RBM microchips. Moreover, we demonstrated a platform for modeling physiological tissue environment on-chip that also allows for rapid cell extraction and cell separation, which is also simple to use and does not require specialized materials.

This reversible bonding approach for microchips is an important addition to the microfluidics technology toolbox, enabling a wider range of on-chip cellular studies. The development of such a versatile, user-friendly approach for reversible bonding fabrication and efficient cell extraction could significantly impact biomedical applications of microfluidics, advancing further development in cell culture or organ-on-chip as well as its following studies such as drug discovery, disease modelling, and cellular interaction analysis.

## Author Contributions

Conceptualization: X.F., X.L., A.R.W.. Methodology: X.F., X.L., L.K.W.C.. Data analysis: X.F., X.L.. Supervision: X.L., A.R.W.. Writing – original draft: X.F., X.L., A.R.W. Writing – review & editing: X.F., X.L., A.R.W..

## Conclusions Conflicts of interest

A.R.W., X.L. and X.F. have filed a USPTO patent on the Reversibly-bonded Microfluidic Devices, application number 63/591,113, filed 17 October 2023. All other authors declare that they have no competing interests.

## Funding

Hong Kong Research Grants Council General Research Fund 16209820 (AW). Center for Aging Science, The Hong Kong University of Science and Technology, Z1003 (AW). Innovation and Technology Commission ITCPD/17-9 (AW).

We would like to acknowledge the Nanosystem Fabrication Facility (CWB) of the HKUST for the device/system fabrication; We would like to acknowledge the technical support from SUSTech; SEM data were obtained using equipment maintained by the Center for Engineering Material and Reliability of Guangzhou HKUST Fok Ying Tung Research Institute, with assistance of Mr. Zhenyu Pan.

For helpful discussion and suggestions, we thank Prof. Rio Ryohichi SUGIMURA at Hong Kong University and his lab student Yang Xiang. We also thank all the members of the A. R. Wu research group for helpful discussions and administrative assistance during this project.

